# Application of a ^1^H Brain MRS Benchmark Dataset to Deep Learning for Out-of-Voxel Artifacts

**DOI:** 10.1101/2023.05.08.539813

**Authors:** Aaron T. Gudmundson, Christopher W. Davies-Jenkins, İpek Özdemir, Saipavitra Murali-Manohar, Helge J. Zöllner, Yulu Song, Kathleen E. Hupfeld, Alfons Schnitzler, Georg Oeltzschner, Craig E. L. Stark, Richard A. E. Edden

**Author notes:** Corresponding Author: Richard A. E. Edden, Ph.D. Department of Radiology and Radiological Science, Johns Hopkins School of Medicine 600 N. Wolfe St, Park 338, Baltimore, MD 21287.

## Abstract

Neural networks are potentially valuable for many of the challenges associated with MRS data. The purpose of this manuscript is to describe the AGNOSTIC dataset, which contains 259,200 synthetic ^1^H MRS examples for training and testing neural networks. AGNOSTIC was created using 270 basis sets that were simulated across 18 field strengths and 15 echo times. The synthetic examples were produced to resemble *in vivo* brain data with combinations of metabolite, macromolecule, residual water signals, and noise. To demonstrate the utility, we apply AGNOSTIC to train two Convolutional Neural Networks (CNNs) to address out-of-voxel (OOV) echoes. A Detection Network was trained to identify the point-wise presence of OOV echoes, providing proof of concept for real-time detection. A Prediction Network was trained to reconstruct OOV echoes, allowing subtraction during post-processing. Complex OOV signals were mixed into 85% of synthetic examples to train two separate CNNs for the detection and prediction of OOV signals. AGNOSTIC is available through Dryad and all Python 3 code is available through GitHub. The Detection network was shown to perform well, identifying 95% of OOV echoes. Traditional modeling of these detected OOV signals was evaluated and may prove to be an effective method during linear-combination modeling. The Prediction Network greatly reduces OOV echoes within FIDs and achieved a median log_10_ normed-MSE of –1.79, an improvement of almost two orders of magnitude.

## 1. Introduction

Proton (^1^H) magnetic resonance spectroscopy (MRS) non-invasively measures levels of endogenous neurometabolites. MRS-visible metabolites are present at millimolar concentrations in the brain, yielding detectable signals with relatively low signal-to-noise ratio (SNR) which mutually overlap. *In vivo* spectra suffer from several artifacts that complicate modeling and interpretation of the data, including eddy current effects and out-of-voxel (OOV) echoes (Kreis, 2004). While there is some degree of standardization and consensus around pre-processing, modeling, and quantification of MRS data (Maudsley et al., 2021; Near et al., 2021; Öz et al., 2021; Wilson et al., 2019), this is an evolving field without a single ideal solution due to the complexity of the problem, and which therefore is likely to benefit from recent advances in machine learning.

Deep learning (DL) uses a network consisting of a series of computational layers to process information (Lecun et al., 2015). Iterative training allows features of the data to be identified and weighted to estimate a final function which predicts a desired output based on a given input (Goodfellow et al., 2016). Supervised learning involves training the network based on a pre-defined target, associating ground-truth parameters with each input. An extensive, balanced, and diverse dataset is preferred to increase the generalizability of the DL outcome. High-dimensional data, such as medical images or time series, are demonstrated to be the most beneficial set of data for several computer vision tasks, such as classification, registration, segmentation, reconstruction, and object detection (Gassenmaier, Küstner, et al., 2021; Lundervold & Lundervold, 2019).

DL has been developed for MRS data as a proof-of-concept in many applications, including metabolite quantification (Chandler et al., 2019; Hatami et al., 2018; H. H. Lee & Kim, 2019, 2020; Rizzo et al., 2023; Shamaei et al., 2023; Zhang & Shen, 2023), signal separation (Li et al., 2020), phase and frequency correction (Ma et al., 2022; Shamaei et al., 2023; Tapper et al., 2021), reconstruction of missing data (H. Lee et al., 2020), accelerated post-processing (Gurbani et al., 2019; Iqbal et al., 2021), denoising (Chen et al., 2023; Dziadosz et al., 2023; Lam et al., 2020), super-resolution (Gassenmaier, Afat, et al., 2021; Iqbal et al., 2019), artifact removal (Gurbani et al., 2018; Kyathanahally et al., 2018), and anomaly detection (Jang et al., 2021). Despite the potential, these methods have yet to be shown to generalize outside of small datasets with a single fixed acquisition protocol. Whereas ‘classical’ methods for post-processing are often driven by an understanding of the problem to be solved, and therefore can often be applied broadly, deep learning methods cannot be assumed to function well outside of the specific datasets used for training and testing. Broadly applicable deep learning methods will only arise from broad training and testing. A key barrier is the lack of a generalized benchmark dataset for training and testing, to play the role that MNIST, ImageNet, and COCO have played in the field of Computer Vision (Fei-Fei et al., 2010; Li Deng, 2012; T.-Y. Lin et al., 2014). Such a dataset lowers the barrier to entry for neural network development in MRS and facilitates performance comparisons between models. The Synthetic Data Working Group of the MRS study group of the International Society for Magnetic Resonance in Medicine’s Synthetic Data Working Group has recently highlighted the MRS community’s need for such a resource. The ultimate goal of this work is to bridge the gap from the synthetic to the in vivo domain, including the additional domain-shift to clinical data.

OOV echoes, which represent a subset of the artifacts often referred to as ‘spurious’ or ‘ghost’ echoes (Kreis, 2004), are a substantial issue for *in vivo* MRS, and an under-studied potential DL application. MRS voxel localization is achieved via a combination of RF pulses and magnetic field gradients, with the intended coherence transfer pathway selected both by phase cycling and dephasing “crusher” gradient scheme (Bodenhausen, 2011). OOV signals arise from gradient echoes – signals from outside the shimmed voxel of interest are refocused by evolution in local field gradients that are either inherent (from air-tissue-bone interfaces) or arising from second-order shim terms (Starck et al., 2009). Therefore, brain regions close to air cavities (e.g., medial prefrontal cortex) or which require stronger shim gradients (e.g., thalamus, hippocampus, etc.) most commonly exhibit OOV artifacts (Starck et al., 2009). OOV echoes seldom occur at the time of the primary echo, so they manifest in the spectrum as broad peaks with strong first-order phase “ripple” that can obscure metabolite resonances. While acquisition strategies can mitigate OOV echoes to some extent, by careful consideration of crusher schemes or voxel orientation (Ernst & Chang, 1996; Landheer & Juchem, 2019; Song et al., 2023), post-processing strategies remain valuable where complete elimination is not possible.

This manuscript develops **A**daptable **G**eneralized **N**eural-Network **O**pen-source **S**pectroscopy **T**raining dataset of **I**ndividual **C**omponents (AGNOSTIC), a dataset consisting of 259,200 synthetic MRS examples. AGNOSTIC spans a range of field strengths, echo times, and clinical profiles, representing metabolite signals, macromolecule (MM) background signals, residual water signals, and Gaussian noise as separate components. To date, DL applications to MRS have relied upon narrow in-house-generated training datasets that limit the generalizability of the solutions developed and comparisons between tools; AGNOSTIC is proposed as a benchmark dataset to fill this gap. In order to demonstrate the utility of this resource, we then illustrate a specific augmentation of the AGNOSTIC dataset to train neural networks for the detection and prediction of OOV echoes.

## 2. Methods

### 2.1. AGNOSTIC Synthetic Dataset

The parameter space that AGNOSTIC spans is deliberately broad, comprising: 18 field strengths; 15 echo times; broad distributions of metabolite, MM, and water amplitudes; and densely sampled time-domain to allow down-sampling. Calculations were carried out using an in-house and open-source Python 3 (Van Rossum & Drake, 2009) programming script using NumPy (Harris et al., 2020). The decision to use in-house software was motivated by needing the flexibility to simulate basis sets that could be manipulated on a spin-by-spin basis which could, for instance, allow for different spins within the same metabolite to have different relaxation rates (e.g., Cr_3.9_ and Cr_3.0_). The dataset is structured as a zipped NumPy archive file (.npz) and can be opened as a Python 3 dictionary object. This zipped NumPy file contains complex-valued NumPy arrays of time-domain (4096 timepoints) data corresponding to the metabolite, macromolecule, water, and noise components which can be combined in different ways depending on the application. For instance, a denoising model may want to target the combined metabolite, MM, and water signal without noise. Within the file, all the acquisition parameters (field strength, echo time, spectral width, etc.), simulation parameters (signal to noise, full-width half-max, concentrations, T_2_ relaxation, etc.), and data augmentation options are specified as detailed below.

#### 2.1.1. Basis Set Simulation

Metabolite spectra are based upon density-matrix-simulated basis functions (Blum, 1981; Fano, 1957; Farrar, 1990; O. W. Sørensen et al., 1984). A total of 270 basis sets were created across 18 field strengths (1.4 T – 3.1 T in steps of 0.1 T) and 15 echo times (10 ms – 80 ms in steps of 5 ms). The Point RESolved Spectroscopy (PRESS) pulse sequence (Bottomley, 1982) was simulated using ideal pulses with TE1 = TE2. The simulated “acquisition window” was started immediately after the last pulse to generate points before the echo. Each metabolite basis was output as an N x 16684 NumPy array, where N is the number of spins for a given metabolite and 16684 is the fixed length of complex time points (300 points before the echo maximum, with an appropriate padding number of zeros and followed by the simulated pre-echo signal, and 16384 points after the echo). The simulated spectral width, centered on 4.7 ppm, was 63.62 ppm for all field strengths (e.g., 8 kHz at 3 T; 4 kHz at 1.5 T). By subsampling the intentionally long time-domain points in the basis set, we can achieve a series of different spectral widths within the ranges commonly seen for *in vivo* experiments without the need to re-simulate the signal with different dwell times.

39 brain metabolite basis functions were simulated: Adenosine Triphosphate (ATP); Acetate (Ace); Alanine (Ala); Ascorbate (Asc); Aspartate (Asp); β-hydroxybutyrate (bHB); β-hydroxyglutarate (2HG); Citrate (Cit); Cysteine (Cys); Ethanolamine (EA); Ethanol (EtOH); Creatine (Cr); y-Amino-Butyric Acid (GABA); Glucose (Glc); Glutamine (Gln); Glutamate (Glu); Glycerophosphocholine (GPC); Glutathione (GSH); Glycerol (Glyce); Glycine (Gly); Water (H_2_O); Homocarnosine (HCar); Histamine (Hist); Histidine (His); Lactate (Lac); Myo-Inositol (mI); N-Acetyl-Aspartate (NAA); N-Acetyl-Aspartate-Glutamate (NAAG); Phenylalanine (Phenyl); Phosphocholine (PCho); Phosphocreatine (PCr); Phosphoethanolamine (PE); Scyllo-Inositol (sI); Serine (Ser); Taurine (Tau); Threonine (Thr); Tryptophan (Trp); Tyrosine (Tyr); and Valine (Val). GABA was separately simulated using two different spin-system enumerations (Govindaraju et al., 2000; Near et al., 2012). Both α-glucose and β-glucose were simulated.

#### 2.1.2. Assembly of Metabolite Component

Individual metabolite basis functions were linearly combined to give a metabolite spectral component, weighted by metabolite concentrations sampled from distributions defined by our recent meta-analysis (Gudmundson et al., 2023), including both healthy and clinical cohort ranges. From the full basis sets, 22 metabolites were selected which had defined concentration ranges available in a recent meta-analysis that collated results from nearly 500 MRS papers using the Preferred Reporting Items for Systematic Reviews and Meta-Analyses (Gudmundson et al., 2023; Moher et al., 2009; Page et al., 2021). One isomer of GABA (either the definition from (Govindaraju et al., 2000) or (Near et al., 2012)) and Glucose (α or β) were randomly chosen with equal probability for each example. Concentrations were selected with equal probability from a range defined by ±2.5 standard deviations from the meta-analysis mean of each cohort (Gudmundson et al., 2023) and are provided in supplemental tables 1 and 2.

T_2_* relaxation decay of time-domain data was simulated with an exponential and Gaussian component to produce a Voigt lineshape (Marshall et al., 1997) in the frequency domain. The exponential component represents the pure T_2_ arising from dipole-dipole interactions, paramagnetic interaction, etc., while the Gaussian component represents the transverse dephasing from diffusion and exchange of spins in an inhomogeneous field (Koch et al., 2009; Marshall et al., 1997; Michaeli et al., 2002; Yablonskiy & Haacke, 1994). While pure T_2_ is understood to be field-independent (Bloembergen et al., 1948; Carr & Purcell, 1954; Held et al., 1973; Michaeli et al., 2002), the dominant Gaussian decay (Marshall et al., 1997) increases with increasing static field strength and is attributed to greater microscopic (Michaeli et al., 2002) and macroscopic (Juchem et al., 2021; Tkáč et al., 2001) susceptibility gradients. Here, the pure Lorentzian T_2_ component is based upon the relaxation times at 1.5 T from a relaxation meta-regression (Gudmundson et al., 2023), which are assumed to be the least impacted by susceptibility gradients that scale with B_0_ (Bloembergen et al., 1948; De Graaf et al., 2006; Michaeli et al., 2002). Once the Lorentzian T_2_ component was applied, the additional T_2_* contributions were modeled by applying appropriate amounts of Gaussian broadening, to achieve a frequency-domain full-width half-maximum (*FWHM*) linewidth of the NAA singlet between 3 Hz and 18 Hz with a uniform distribution. A small amount of jitter (between 20 s^-2^ and 100 s^-2^) was added to the Gaussian decay rate so that each metabolite would undergo a similar, but not identical, amount of Gaussian decay to better replicate the variability observed for *in vivo* data.

#### 2.1.3. Macromolecular Component

Fourteen MM signals were modeled at: 0.92 ppm; 1.21 ppm; 1.39 ppm; 1.67 ppm; 2.04 ppm; 2.26 ppm; 2.56 ppm; 2.70 ppm; 2.99 ppm; 3.21 ppm; 3.62 ppm; 3.75 ppm; 3.86 ppm; and 4.03 ppm (Cudalbu et al., 2021; Giapitzakis et al., 2018). MM chemical shifts were jittered by ± 0.03 ppm to both account for observed differences in MM designations and provide further dataset augmentation. Each MM signal was simulated as a singlet with exponential decay rate sampled uniformly from a range specified by literature of MM T_2_ time constants (Murali-Manohar et al., 2020) and additional Gaussian decay to reach published linewidths (Giapitzakis et al., 2018; Murali-Manohar et al., 2020). MM amplitudes were sampled uniformly from within published ranges (Giapitzakis et al., 2018; Murali-Manohar et al., 2020).

#### 2.1.4. Noise Component

Noise was generated from a normal distribution, with independent random real and imaginary points. The noise was scaled such that the signal-to-noise ratio of the NAA singlet (SNR_NAA_ was defined, following Experts’ Consensus (Öz et al., 2021), by *NAA height divided by the standard deviation of the noise*) was uniformly sampled between 5 and 80. The noise amplitude values are also stored within the archive file.

#### 2.1.5. Residual Water Component

The residual water basis signal was simulated as a composite signal (of up to five components). In order to simulate varying degrees of water suppression, the residual water signal was modeled by between 0 and 5 unique Voigt-shaped signals with variable ppm locations, phases, and amplitudes, based on the approach of (L. Lin et al., 2019). The ranges for these parameters are listed in Table 1. The final water component was scaled to be between 1× and 20× the maximum value of the frequency-domain metabolite spectrum. The water components used, along with their corresponding parametrizations, are stored within the NumPy archive file.

**Table 1.** Parametrization of the residual water signal components within AGNOSTIC.

| Component | Location / ppm |  | Phase / deg |  | Amplitude |  |
| --- | --- | --- | --- | --- | --- | --- |
|  | Low | High | Low | High | Low | High |
| 1 | 4.679 | 4.711 | −10 | 10 | 1.00 | 1.00 |
| 2 | 4.599 | 4.641 | 15 | 45 | .35 | .55 |
| 3 | 4.759 | 4.801 | −60 | −30 | .35 | .55 |
| 4 | 4.449 | 4.541 | −70 | 45 | .10 | .25 |
| 5 | 4.859 | 4.901 | 105 | 135 | .10 | .25 |

#### 2.1.6. Frequency and Phase Shifts

Within the NumPy archive file, a frequency shift, zero-order phase shift, and first-order phase shift are specified for each entry in the dataset, but not applied to the time-domain components. Frequency shifts were sampled uniformly from the range −0.313 ppm to +0.313 ppm. Zero-order phase shifts were sampled uniformly from the range −180 degrees to +180 degrees. First-order phase shifts were sampled uniformly from the range −19.5 degrees to +19.5 degrees per ppm. Users may choose to omit phase and frequency shifts, use the provided shifts, or specify their own.

### 2.2. Exemplar Application to AGNOSTIC: Machine Learning for Out-Of-Voxel Artifacts

The primary motivation for the AGNOSTIC dataset is as a training resource for the development of processing, modeling, and analysis tools for MRS. Synthetic spectra with known ground truths are valuable in a range of applications, from the development and validation of traditional linear combination modeling algorithms to training DL models.

In order to demonstrate the utility of the dataset, an exemplar application is presented, in which the AGNOSTIC dataset is supplemented by simulated artifacts (in this case out-of-voxel OOV echoes) and used to train DL models to detect and predict the artifact signals. The AGNOSTIC dataset was developed as building blocks which can be combined to train a variety of different models. A strength of this dataset is that custom user-defined components can be utilized. We demonstrate this point here by building an OOV dataset to train and evaluate a DL model to identify and suppress OOV artifacts.

#### 2.2.1. Simulation of Out-Of-Voxel Echoes

OOV artifacts were defined as complex time-domain signals with a time point (τ_OOV_), width (W_OOV_), frequency (ω_OOV_), phase (Φ_OOV_) and amplitude (a_OOV_) as shown in Figure 1. τ_OOV_ describes the timepoint of the top of the OOV echo and was sampled randomly from a uniform distribution between 10 ms and 400 ms. W_OOV_ describes the Gaussian decay rate and was sampled randomly from a uniform distribution between 500 s^-2^ and 8000 s^-2^, resulting in a FWHM echo duration between 18 ms and 74 ms. ω_OOV_ describes the offset in the frequency domain, and was sampled randomly from a uniform distribution in order to produce OOVs that occur between 1 ppm and 4 ppm. a_OOV_ was sampled randomly from a uniform distribution to produce OOV echoes with an amplitude between 0.1% and 20% of the maximum time domain point. Φ_OOV_ was sampled uniformly between 0 degrees to 360 degrees.

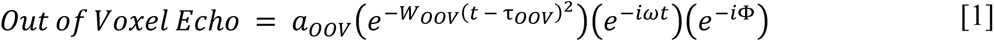

**Figure 1.**
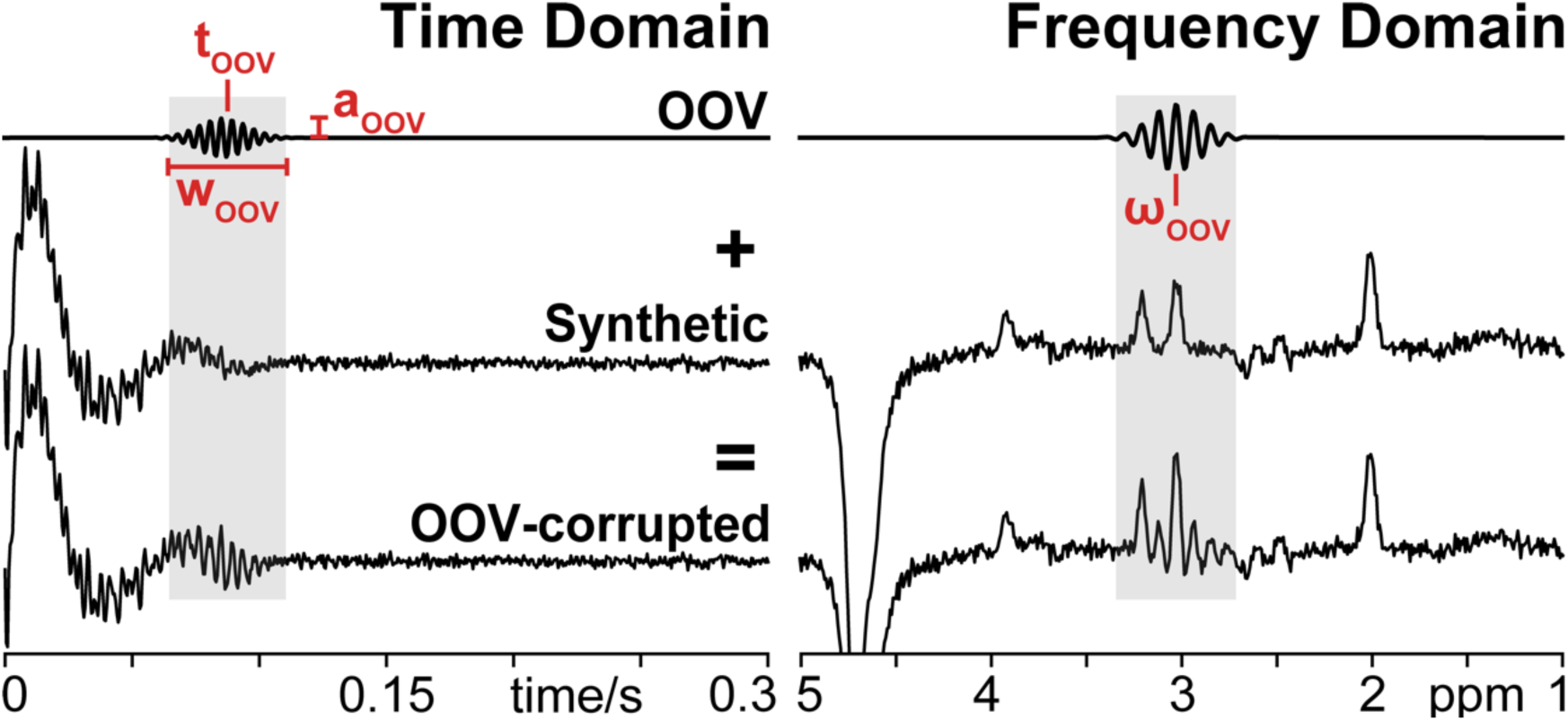
Simulation of OOV echoes and OOV-corrupted synthetic data: OOV echoes were simulated as complex time-domain signals with a center timepoint (τ_OOV_), width (W_OOV_), frequency (ω_OOV_), phase (Φ_OOV_), amplitude (a_OOV_). OOV echoes were added to 85% of synthetic data to create datasets for training and evaluation.

#### 2.2.2. Integration of OOV Echoes into AGNOSTIC for the Training Dataset

To build the OOV echoes dataset, we combined metabolite, water, MM, and noise components from the AGNOSTIC dataset. We then added OOV signals to 85% of the dataset and a complex zeros array in the remaining 15%. In total there were 180,000 examples used for network training, 1,800 examples used for validation, and 7,200 examples used for testing. Finally, we applied the included frequency and phase shifts specified within the AGNOSTIC dataset. The network input consisted of the combined metabolite, water, MM, noise, and OOV signals as a complex time-domain signal. This input was normalized so that the absolute maximum among the real and imaginary values was 1. Finally, training data were converted to a TensorFlow Dataset (Abadi et al., 2015).

#### 2.2.3. Detection Network

The first exemplar network is designed to detect OOV echoes within time-domain data by identifying the points in the spectra that have been contaminated by OOV echoes. This Detection Network is a fully Convolutional Neural Network (CNN) designed using TensorFlow2 with Keras (Chollet & others, 2015) in a Python 3 environment. The network consists of contracting encoding layers and expanding decoding layers with a total of 1.543 million parameters, as shown in Figure 2. Each layer was initialized (kernel_initializer) with “he_normal” (He et al., 2015). Each convolutional layer (except the output layer which uses a sigmoid activation) includes batch normalization and a leaky rectified linear unit (ReLu) activation function (Maas et al., 2013)Click or tap here to enter text.. A kernel size of 3 (3 x 2 before collapsing the real/imaginary dimension and 3 x 1 afterward) was used for each convolutional layer. The network is designed to receive a time-domain input signal and return a binary mask of the same size as the input with ones placed in OOV-detected regions and zeros elsewhere. A ground-truth binary mask was determined as the 5% level of the maximum amplitude of the Gaussian OOV envelope located at the central peak. For training, the input and output of this network is a 60 x 2048 x 2 x 1 tensor, where 60 is the batch size, 2048 is the number of time points, 2 is the real/imaginary dimension, and 1 is the channel dimension.

**Figure 2.**
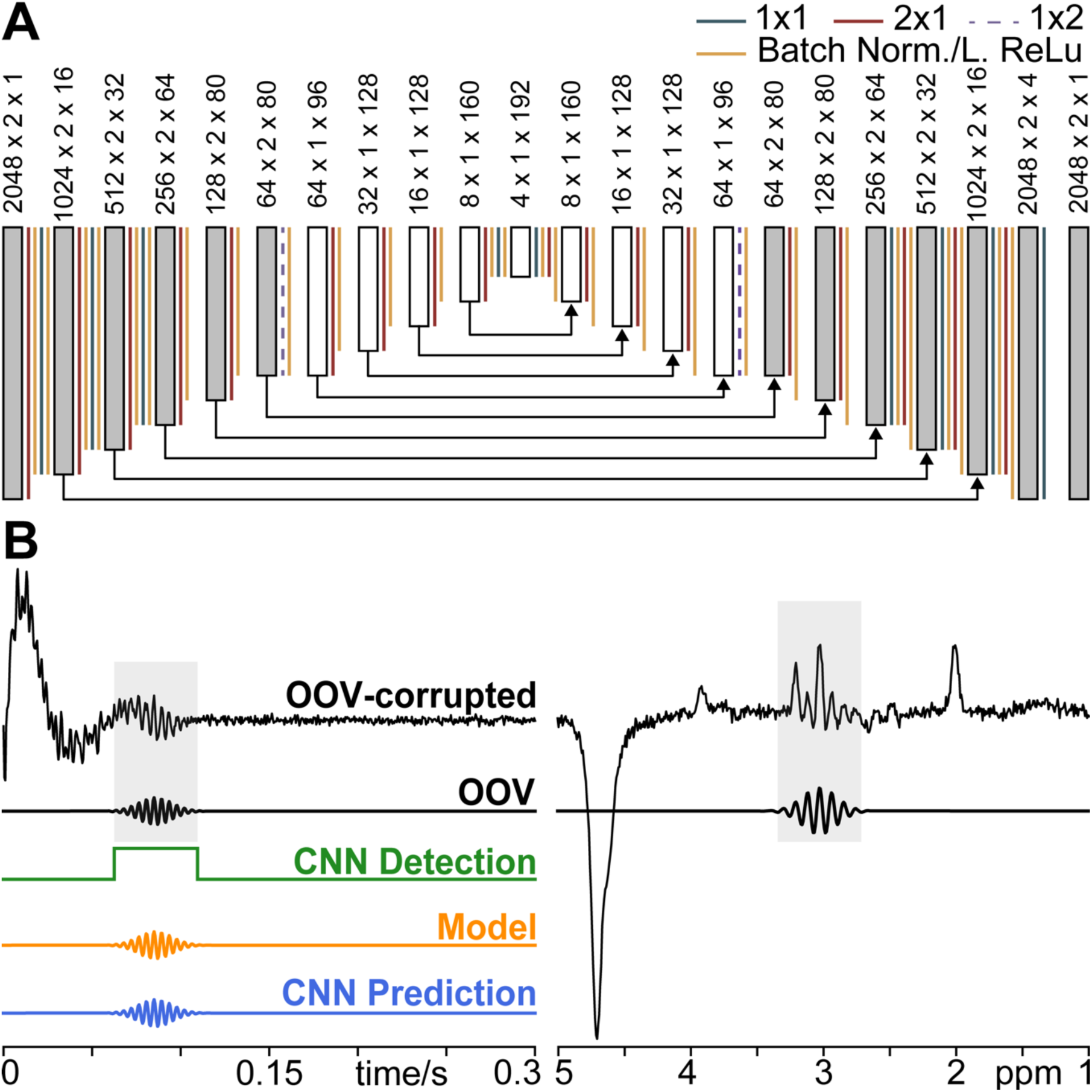
Convolutional Neural Network Architecture, Input, and Output: A) Fully convolutional neural network architecture used for both the Detection and Prediction Network. Convolutional strides, batch normalization, and Leaky ReLu activation functions are denoted by a colored line. Dark gray blocks represent complex data with the 2nd dimension representing real and imaginary components, while white blocks represent the network abstracted single dimension. Arrows show residual connections. Note, inputs and outputs are all time-domain signals; Frequency-domain is shown for convenient visualization. B) OOV-corrupted synthetic example and the isolated OOV. The complex OOV-corrupted data was used as the Detection and Prediction Network input. The target Output is the isolated OOV.

The Dice coefficient (Carass et al., 2020; Dice, 1945; T. Sørensen, 1948) of the overlap between the network output and the correct binary OOV location vector was used as a training loss function, calculated as 2x the intersection divided by the union plus 1; where 1 was used to avoid division by 0. The Adam (Kingma & Ba, 2015) optimizer was used with a fixed learning rate of 0.0003. Success on the validation set was evaluated every 7,200 steps, at which time the network weights were saved if the validation loss improved. The final model that was selected had the smallest validation loss after 72 epochs. Training took approximately 2.5 hours and was performed on an 8 GB NVIDIA GeForce RTX 3070 GPU. A clustering algorithm was applied to the final network output, which zeroed any group of time points in which the network detected OOV echo that was smaller than 5 consecutive time points, to dampen spurious output. A cluster size of 5 was selected empirically to ensure detection of the narrow echoes while eliminating any false positives.

#### 2.2.4. Modeling

Modeling of the OOV echoes was performed as an optimization problem and solved with SciPy (Virtanen et al., 2020) minimization routines. Here, the non-gradient Powell (Powell, 1964, 1994) optimizer was used to determine the five OOV parameters (τ_OOV_, W_OOV_, ω_OOV_, Φ_OOV_, and a_OOV_), minimizing the mean squared error (MSE) between the model and the data within the time window identified by the Detection Network. Initial values for τ_OOV_, W_OOV_, and the a_OOV_ are inferred from the Detection Network’s output center timepoint, the detection duration, and the standard deviation of the target signal within the detected region.

Optimization was performed as three sequential optimization steps performed one after another. The first optimization is used to determine τ_OOV_, W_OOV_, and the a_OOV_ by minimizing the MSE between the absolute values of the model and the data (i.e., removing frequency and phase from the model) in the time domain. The second optimization determines ω_OOV_ by minimizing between the absolute values of the model and the data in the frequency domain. The third optimization refines the values determined by optimizations 1 and 2 and determines Φ_OOV_ by complex optimization in the time domain.

#### 2.2.5. Prediction Network

The second exemplar network is designed to predict the OOV echoes found within time-domain data. This prediction network is also a fully CNN designed using TensorFlow2 with Keras in a Python 3 environment, with the same architecture as the Detection Network (as shown in Figure 2). As such, the input and output of this network is also 60 x 2048 x 2 x 1 tensor, where 60 is the batch size, 2048 is the number of time points, 2 is the real/imaginary dimension, and 1 is the channel dimension. The network is designed to receive a time-domain input signal containing a combination of the ground-truth time-domain signal and the OOV artifact and return a time-domain output signal that only contains the OOV signal, amplified 10x. This amplification serves to focus the training on the OOV echo by non-uniformly (due to the OOV echo’s non-linear decay) concentrating the network towards the center-most points of the OOV echoes to effectively center and reconstruct the predicted OOV echo on the τ_OOV_ with the correct W_OOV_.

For training, a weighted mean squared error (weighting the timepoints within the ground-truth OOV mask uniformly by 10) was used as a loss function with an ADAM (Kingma & Ba, 2015) optimizer and a fixed learning rate of 0.0003. Success on the validation set was evaluated every 7,200 steps at which time the network weights were saved if the validation loss improved. The final model that was selected had the smallest validation loss after 72 epochs. Training took approximately 2.5 hours and was performed on an 8 GB NVIDIA GeForce RTX 3070 GPU.

#### 2.2.6 Evaluating the Performance of Networks and Modeling

In the final testing set, OOV artifacts were present in 6,137 of the total 7,200 examples (85.2%). The Detection Network was evaluated using the Dice coefficient (Carass et al., 2020; Dice, 1945; Powell, 1964), the overlap between the ground-truth binary OOV mask and the cluster-thresholded network output. As well as computing global success, the dependence of detection success on various attributes of the OOV echo and the underlying spectrum were also investigated.

Both modeling and the prediction network return a pure OOV signal, and in both cases, the MSE between the prediction/model and the ground-truth OOV echo is used for evaluation. If the ground-truth echo datapoints are E_i_ and the model or echo prediction is M_i_, we calculate the fractional remaining OOV amplitude as:

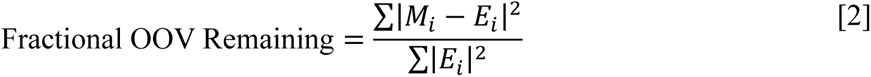

where the bars represent the complex amplitude. The sum is taken over the ground-truth range of the OOV echo. In order to visualize a wide range of success and failure, we take the log_10_ of this quantity for plotting (i.e., a log_10_ value of 1 is no change, anything positive is a manipulation that is worse than doing nothing, and a negative value show the order of magnitude of improvement). Note that *E_i_* is the ground-truth echo signal, not the signal from which the echo is being removed which also contains metabolite, macromolecule, and noise components.

The timing of the OOV was found to be a key parameter determining the success of detection and prediction, and as a result, the evaluation metrics were calculated for the following time-bins (based on the known value of t_OOV_): 10-20 ms; 20-40 ms; 40-60 ms; 60-80 ms; 80-120 ms; 120-200 ms; 200-300 ms; 300-400 ms.

#### 2.2.7 *In Vivo* Proof-of-Principle

As a proof-of-principle demonstration of this exemplar use of the AGNOSTIC dataset, the network was applied to 256 transients of *in vivo* data, selected because they contain prominent OOV echoes and were excluded during quality assessment in a recent study (Zöllner et al., 2023). These data were collected on a 2.89 T Siemens scanner using the MEGA-PRESS (Mescher et al., 1996, 1998) pulse sequence with a TE of 68 ms and TR of 1.75 s, and a spectral width of 2.4 kHz. Note that this challenges the generality of the training because the network has never seen data acquired at 2.89 T, nor at 2.4 kHz spectral width, nor at TE 68 ms, nor with MEGA-Editing, nor with actual real RF pulses. Raw data from a 25 x 25 x 25 mm^3^ voxel in the cerebellum were loaded and coil combined in Osprey (Oeltzschner et al., 2020). Time-domain data were saved as a MATLAB (The MathWorks Inc., 2022) .mat file and loaded as a Python 3 object using SciPy. The data were normalized (as above with training data) to be used as input for the neural networks.

One challenge of *in vivo* data (and the reason that this network demonstration focuses substantially on synthetic data) is that no ground truth is available. Therefore, the degree of success in removing OOV echo signals from time-domain data *D_i_* is:

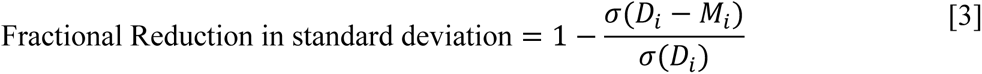

where s denotes the standard deviation. Note that, in contrast to the metric used for synthetic data in Equation 1, only *D_i_* is available, not the ground truth *E_i_*, which substantially changes the ceiling of success. It is still expected that substantial signal variance remains after OOV removal, since *D_i_* contains metabolite signals and noise. The range over which this standard deviation is calculated is the 50% level of the normalized histogram of the detection network’s output across the 256 transients. Note that this metric is an imperfect response to the absence of ground-truth knowledge for in vivo data, predicated on the assumption that subtracting out OOV signal reduces the standard deviation of the time-domain signal.

## 3. Results

### 3.1. AGNOSTIC Synthetic Dataset

The AGNOSTIC dataset contains 259,200 examples, consisting of 960 examples from each of the eighteen field strengths and fifteen echo times (i.e., 960x18x15=259,200). A representative set of ten spectra are shown in Figure 3, illustrating the diversity of field strengths, TEs, SNR, and linewidth within the dataset.

**Figure 3.**
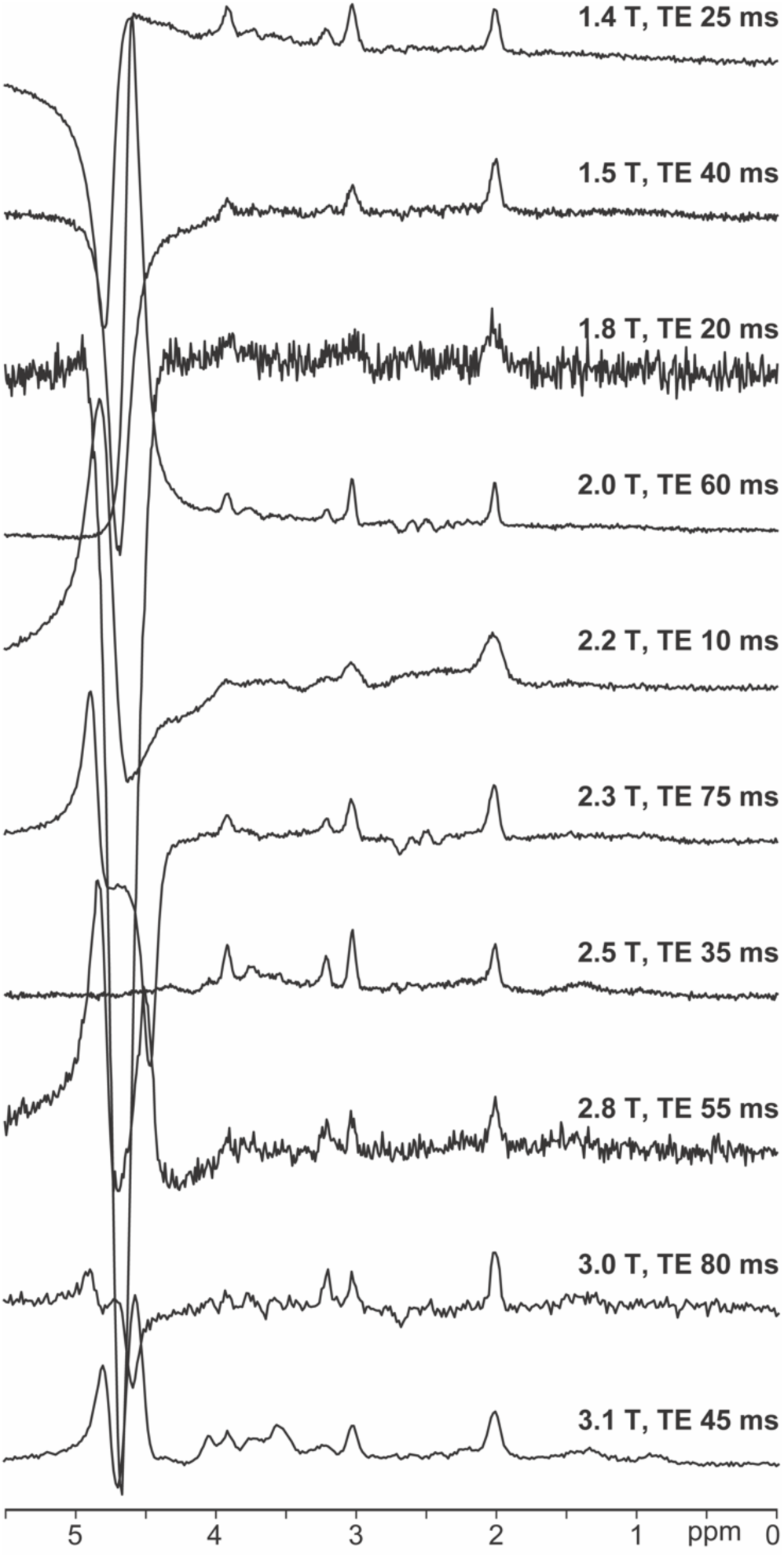
AGNOSTIC synthetic dataset. 10 representative spectra from the AGNOSTIC dataset. The 10 examples show the diversity of field strength, TE, linewidths, and residual water signal present among the data. Note, examples are shown here in the frequency-domain to better illustrate the heterogeneity, but the dataset provides time-domain examples.

One challenge with making this dataset available is its size — 75 GB — but we do make it freely available on Dryad (DOI: 10.7280/D1RX1T). The basis sets from which these are constructed are more manageable — 9 GB — and can also be accessed through Dryad (DOI: 10.7280/D1RX1T). Code for generating the AGNOSTIC dataset locally is available at: https://github.com/agudmundson/agnostic.

### 3.2. Exemplar Application to AGNOSTIC: Machine Learning for Out-Of-Voxel Artifacts

#### 3.2.1. Detection Network

Of the 6,137 examples where OOV artifacts were present, the Detection Network correctly identified 5,827 (94.9%) with a median Dice score of 0.974 (0.941–0.985 interquartile range) and missed 310 (5.05%) with a Dice score of 0.00. In the 1063 examples that did not include OOV artifacts, the network correctly ignored 912 (85.8%) and falsely detected OOV echoes in 151 (14.2%). Figure 4 shows the Detection Network’s output for a synthetic OOV-corrupted example.

**Figure 4.**
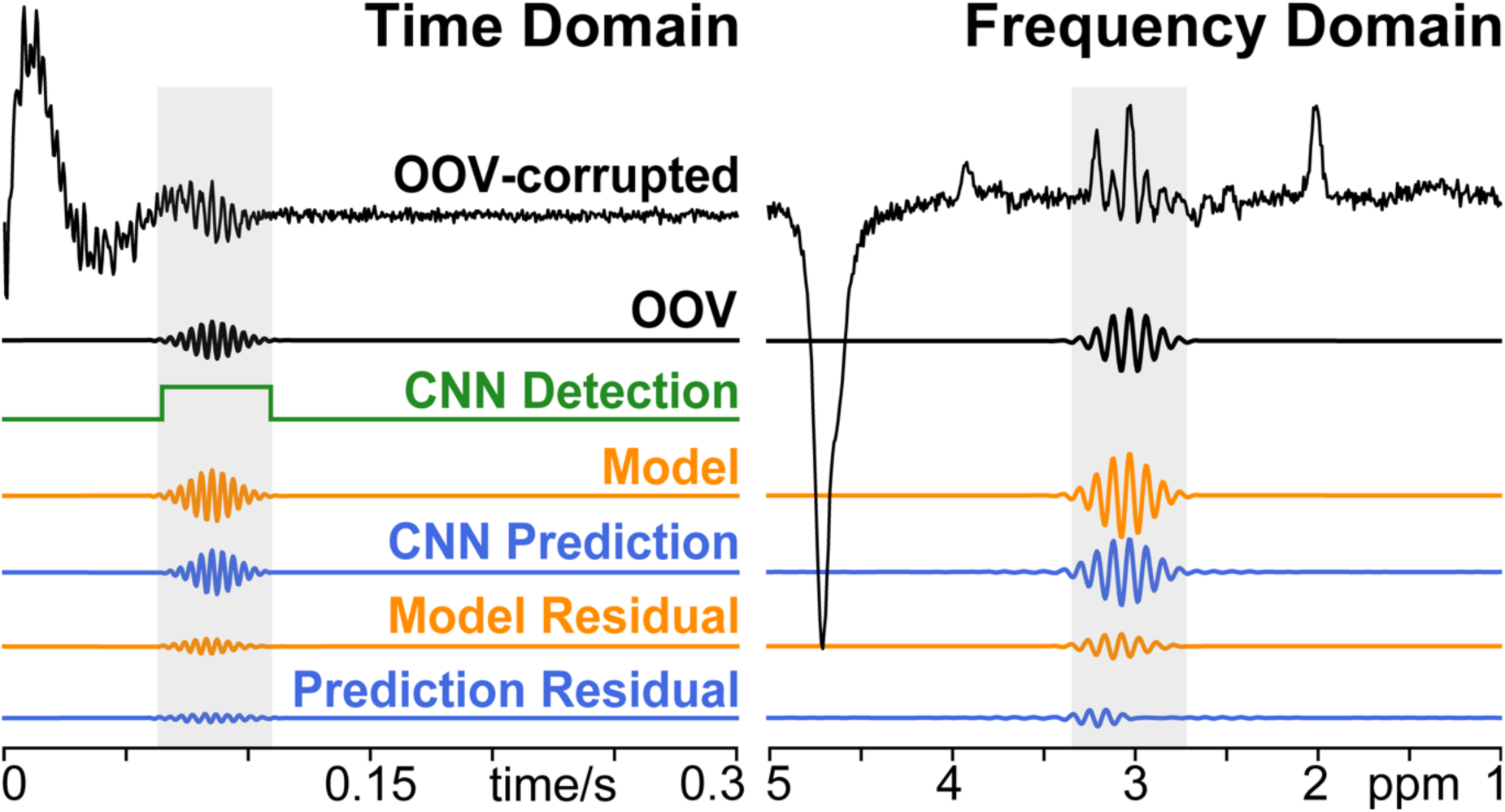
OOV-corrupted example: OOV-corrupted synthetic example and the isolated OOV. Results from Detection Network (green), Model (orange), and Prediction Network (blue) are shown below the ground truth OOV-corrupted and OOV. OOV residuals are shown for the Model (orange) and Prediction Network (blue) demonstrating remaining signal after subtraction. Note, frequency-domain is shown for convenient visualization, but the Detection Network, Modeling, and Prediction Network all operate on time-domain signals.

Analysis of the factors that determined success indicated that the time at which OOV signals occur is most critical. Therefore, OOV echoes were further broken down into eight time-bins, and the Dice score plotted in Figure 5. The median Dice scores — 0.165, 0.858, 0.892, 0.934, 0.960, 0.974, 0.978, and 0.978 — are poor in the first bin and improve thereafter. Note that these bins are not spaced equally to emphasize poor performance extremely early. The number of examples in each bin is 161, 289, 282, 329, 622, 1256, 1565, and 1633, respectively.

**Figure 5.**
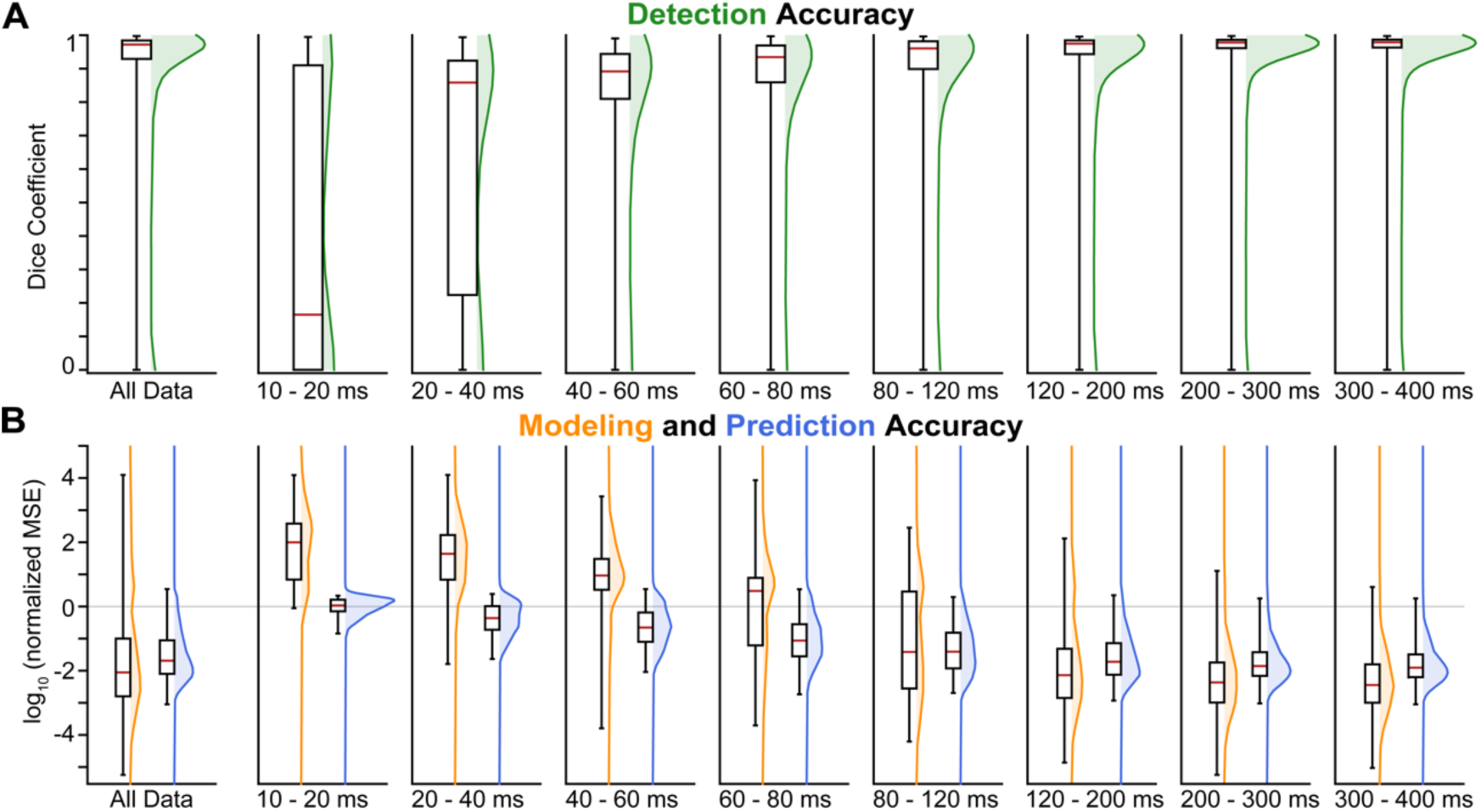
Evaluation of Detection Network, Modeling, and Prediction Network. A testing set with 7200 (2400 examples with 3 different OOV echoes) unseen examples was used to evaluate the A) Detection Network and B) Modeling and Prediction Network. Performance across the whole test set is shown on the left-hand side. Performance across the binned center timepoint (τOOV) is shown across the right-hand side.

#### 3.2.2. Modeling

The modeling optimization converged in 5,824 of the 5,827 examples where the detection network detected OOV artifacts and provided initial values. Across this subset of the examples, the modeling achieved a median log_10_ (fractional OOV remaining) of −2.19 (−2.90 – −1.19 inter-quartile range), i.e., a median reduction of more than two orders of magnitude. Figure 5 shows the resulting model for a synthetic OOV-corrupted example.

These values — broken down into 8 time-bins — are shown in Figure 5. The median log_10_(fractional OOV remaining) decreases across the time bins: 1.663, 1.324, 0.680, 0.223, −1.586, −2.276, −2.491, and −2.567.

#### 3.2.3. Prediction Network

In the 6,137 examples where OOV artifacts were present, the prediction network achieved a median log_10_ normed-MSE of −1.79 (−2.21 – −1.11 inter-quartile range). In the 5,824 examples where OOV artifacts were successfully modeled, the prediction network achieved a median log_10_ normed-MSE of −1.85 (−2.24 – −1.24 inter-quartile range). Figure 5 shows the Prediction Network’s output for a synthetic OOV-corrupted example.

OOVs were further broken down into 8 time-bins (Figure 5) early — the number of examples in each bin is 86, 226, 261, 312, 592, 1208, 1538, 1601. The median log_10_(fractional OOV remaining) decreases across the time bins: −0.207, −0.583, −0.862, −1.250, −1.577, −1.878, −2.005, and −2.052.

#### 3.2.4. *In Vivo* Proof-of-Principle

The Detection Network identified an OOV in 243 of 256 transients (94.9%). In these 243 OOV-detected transients, the modeling achieved a median reduction in standard deviation of 71.0 % (60.2 -75.3% inter-quartile range). The Prediction Network achieved a median reduction in standard deviation of 69.65% (66.33 %/72.7 % inter-quartile range) in this subset. In the full set of 256 transients, the Prediction Network achieved a median 69.4 % (65.3 – 72.6 % inter-quartile range) reduction in standard deviation. The standard deviation of the noise floor was found to account for a median of 10.3% (9.35–11.6 % inter-quartile range) of the standard deviation of signal within the time window for the 256 averages. A representative *in vivo* example is shown in Figure 6.

**Figure 6.**
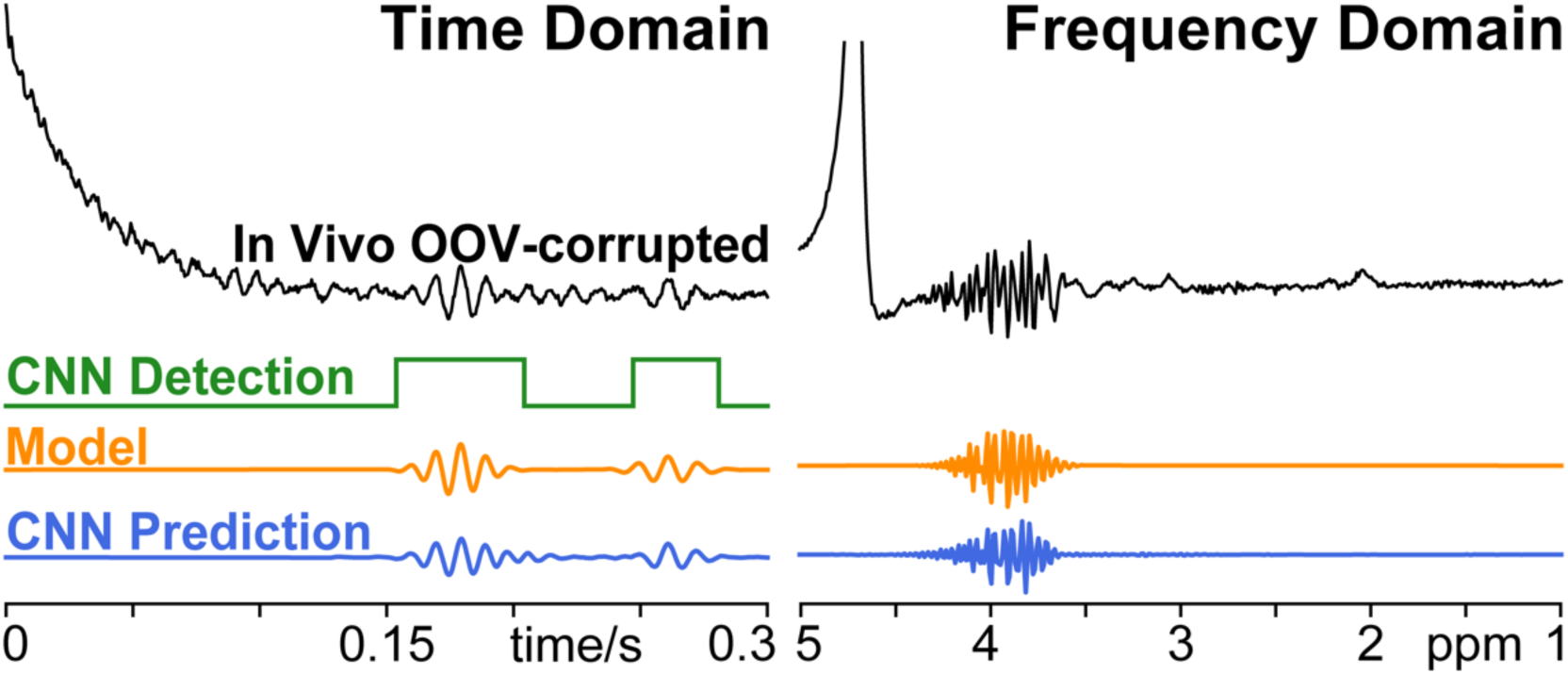
*In vivo* MEGA-PRESS OOV-corrupted example. Results from Detection Network (green), Model (orange), and Prediction Network (blue) are shown below. Detection and Prediction CNNs identified and reconstructed the OOV echo, despite having never seen data acquired with 2.89 T, 2.4 kHz spectral width, 68 ms, editing, nor real RF pulses. Note, frequency-domain is shown for convenient visualization, but the Detection Network, Modeling, and Prediction Network all operate on time-domain signals.

## 4. Discussion

AGNOSTIC is a benchmark MRS dataset for training and evaluating performance across various models. In order to make these synthetic data representative of in vivo brain MRS datasets, a total of 22 brain metabolites and 14 MM peaks were simulated within 270 basis sets, spanning field strengths from 1.4 T to 3.1 T and TEs from 10 to 80 ms. Parameterized water residual and noise were included. SNR and linewidths were assigned at random, independent of B_0_ or TE. The broad span of the dataset is key in training networks that generalize. While AGNOSTIC is broad in these dimensions, it does only represent simulated data for PRESS (Bottomley, 1982) acquisitions, and may benefit from expansion to include other pulse sequences, such as STEAM (Frahm et al., 1987), SPECIAL (Mekle et al., 2009; Mlynarik et al., 2006), LASER (Garwood & DelaBarre, 2001), and semi-LASER (Scheenen, Heerschap, et al., 2008; Scheenen, Klomp, et al., 2008), and edited versions including MEGA (Mescher et al., 1996, 1998) and Hadamard-encoded (Chan et al., 2016, 2019; Oeltzschner et al., 2019; Saleh et al., 2016) schemes. AGNOSTIC is limited by simulations that used ideal pulses, a calculated trade-off to emphasize generalizability across field strength, echo time, and spectral width, and thus fail to capture effects associated with spatially heterogeneous coupling evolution. The extent to which these limitations matter will depend on the applications that AGNOSTIC synthetic data are being used for.

The Detection network was highly successful, identifying 94.9% of the testing set where OOV artifacts were present. The precise value of this success metric is obviously impacted by the parameters of the OOVs – a later minimum OOV time would tend to increase performance, and earlier would degrade it. It is noteworthy that, although the training datasets never contained more than one OOV echo, the detection and prediction networks were able to handle more than one OOV echo in vivo data, presumably because CNNs operate locally within the FID. It is also encouraging that the networks generalized well to the *in vivo* data (Figure 6), which was collected with unseen acquisition parameters, i.e., edited MEGA-PRESS (Mescher et al., 1996, 1998) data acquired at 2.89 T with a TE of 68 ms, and 2.4 kHz spectral width. While it is reasonable to believe that networks trained using AGNOSTIC will generalize well with *in vivo* clinical data, future work will need to evaluate performance for clinical applications.

In the exemplar OOV application, the success of the networks depended heavily on the timing of the OOV signal. The earliest OOV echoes were most challenging, unsurprisingly since such signals are broad Gaussian resonances that are indistinguishable from within-voxel MM and baseline signals. Indeed, the only feature that differentiates OOV signals from other broad components of the model is timing. It is conceptually helpful to consider this in the Fourier domain, even though all network processing is performed in the time domain. In the frequency domain, a mismatch between the echo-top and the acquisition start is represented as a first-order phase error of the signal associated with that echo. Where insufficient first-order phase exists to be represented within the linewidth of the signal in question, (which in the time domain corresponds to substantial truncation of the lefthand side of the echo), the network struggles to identify OOV signals.

In the context of this study, modeling and prediction are treated as two alternative approaches to OOV characterization. For early OOV signals, the modeling approach tended to mis-attribute non-artifact signal as OOV signal, a result that the metric scored as worse than no intervention. The median performance of the Prediction network, even for very early OOV signals, was close to zero. Both modeling and prediction performance improve as the OOV moves later in the acquired signal, with modeling improving faster than the network, and performing better than prediction beyond 120 ms. This strong performance of the model at least in part reflects the exact match between the generative model of the synthetic OOV artifacts and the model that is being used to extract them. More moderate performance might be expected for real in vivo examples – but the same may also be true for networks which have been trained with the same synthetic data and may have learned specifically to identify OOV signals that have a Gaussian kernel.

One key difference between most DL applications and applications in MRS, is the strict requirement to preserve amplitude fidelity in network outputs. A common approach to artifacts in DL is to return an artifact-free version of the network input. In contrast, the approach taken here is to return the artifact, which has the following benefits: it avoids networks over-learning the formulaic pattern of typical spectra; it reduces the impact of the lack of sequence diversity within the AGNOSTIC dataset; and it is less likely to impact the amplitudes of metabolite signals.

The ultimate goal of this work is to extract metabolite levels from MRS data that are not impacted by OOV artifacts. This problem can be addressed at several points: either by not acquiring data that contain OOV artifacts; by removing OOV artifacts post-acquisition; and by incorporating appropriate OOV model components into quantification model so that the impact of OOV is minimized. While the work presented here focuses primarily on the second context, it raises important potential applications in the other contexts. One motivator for developing the Detection network is the possibility of real-time deployment during sequence acquisition to trigger sequence changes when OOV artifacts are detected. The modeling applied here was time-restricted to a given window and ignored other components of the spectrum, but demonstrates potential for future integration within a full linear-combination model.

## 5. Conclusion

In conclusion, we have presented the AGNOSTIC benchmark dataset which can be used for training and testing brain-specific ^1^H MRS deep learning models. This large synthetic dataset is open-source and encompasses a range of field strengths, TEs, and dwell times to ensure networks are robust to a variety of *in vivo* data acquisitions protocols. Using this dataset, we have demonstrated an exemplar use case to develop CNNs to detect and predict out-of-voxel artifacts.

## Abbreviations

^1^H: proton
2HG: β-hydroxyglutarate
Ace: acetate
AGNOSTIC: adaptable generalized neural-network open-source spectroscopy training dataset of individual components
Ala: alanine
Asc: ascorbate
Asp: aspartate
ATP: adenosine triphosphate
bHB: β-hydroxybutyrate
Cho: choline-containing compounds
Cit: citrate
CNN: Convolutional Neural Networks
Cr: creatine
Cys: Cysteine
DL: deep learning
EA: Ethanolamine
EtOH: ethanol
FID: free induction decay
FWHM: full-width half-maximum
GABA: gamma-aminobutyric acid
Glc: glucose
Gln: glutamine
Glu: glutamate
Glx: sum of glutamate and glutamine
Gly: glycine
Glyce: glycerol
GM: gray matter
GPC: glycerophosphocholine
GSH: glutathione
H2O: water
HCar: homocarnosine
Hist: histamine
His: histidine
ISMRM: international society for magnetic resonance in medicine
Lac: lactate
LASER: localization by adiabatic selective refocusing
MEGA: Mescher-Garwood
mI: myo-inositol
MM: macromolecule
MRS: magnetic resonance spectroscopy
MSE: mean-squared error
NAA: N-acetylaspartate
NAAG: N-acetyl-aspartyl-glutamate
OOV: out-of-voxel
PCho: phosphocholine
PCr: phosphocreatine
PE: phosphoethanolamine
Phenyl: phenylalanine
PRESS: point resolved spectroscopy
ReLu: rectified linear unit
Ser: serine
sI: scyllo-inositol
sLASER: semi-adiabatic localization by adiabatic selective refocusing
SPECIAL: spin echo full intensity acquired localiezed
STEAM: stimulated echo acquisition mode
SNR: signal-to-noise ratio
T2: spin-spin relaxation time
Tau: taurine
tCho: sum of choline-containing metabolites
tCr: sum of creatine and phosphocreatine
TE: echo-time
Thr: threonine
tNAA: sum of N-acetyl-aspartate and N-acetyl-aspartyl-glutamate
Trp: Tryptophan
Tyr: Tyrosine
Val: Valine

## Acknowledgments

This work has been supported by The Henry L. Guenther Foundation, Sonderforschungsbereich (SFB) 974 (TP B07) of the German Research foundation, and the National Institute of Health, grants T32 AG00096, R00 AG062230, R21 EB033516, R01 EB016089, R01 EB023963, K00AG068440, P30 AG066519, R21 AG053040, R01 AG076942, P30 AG066519 and P41 EB031771.

## CRediT authorship contribution statement

**Aaron T. Gudmundson:** Conceptualization, Data Curation, Formal Analysis, Investigation, Methodology, Software, Visualization, Writing-original draft, Writing-Review & Editing. **Christopher W. Davies-Jenkins:** Data Curation, Resources, Writing-original draft, Writing-Review & Editing. **İpek Özdemir:** Data Curation, Writing-original draft, Resources, Writing-Review & Editing. **Saipavitra Murali-Manohar:** Data Curation, Writing-original draft, Writing-Review & Editing. **Helge J. Zöllner:** Data Curation, Resources, Writing-original draft, Writing-Review & Editing. **Yulu Song:** Data Curation, Resources, Writing-original draft, Writing-Review & Editing. **Kathleen E. Hupfeld:** Writing-original draft, Writing-Review & Editing. **Alfons Schnitzler:** Data Curation, Resources, Writing-Review & Editing. **Georg Oeltzschner:** Conceptualization, Supervision, Writing-original draft, Writing-Review & Editing. **Craig Stark:** Conceptualization, Funding acquisition, Project Administration, Resources, Supervision, Writing-original draft, Writing-Review & Editing. **Richard A.E. Edden:** Conceptualization, Funding acquisition, Project Administration, Resources, Supervision, Visualization, Writing-original draft, Writing-Review & Editing.

## Supplemental Material

**Supplemental Figure 1:**
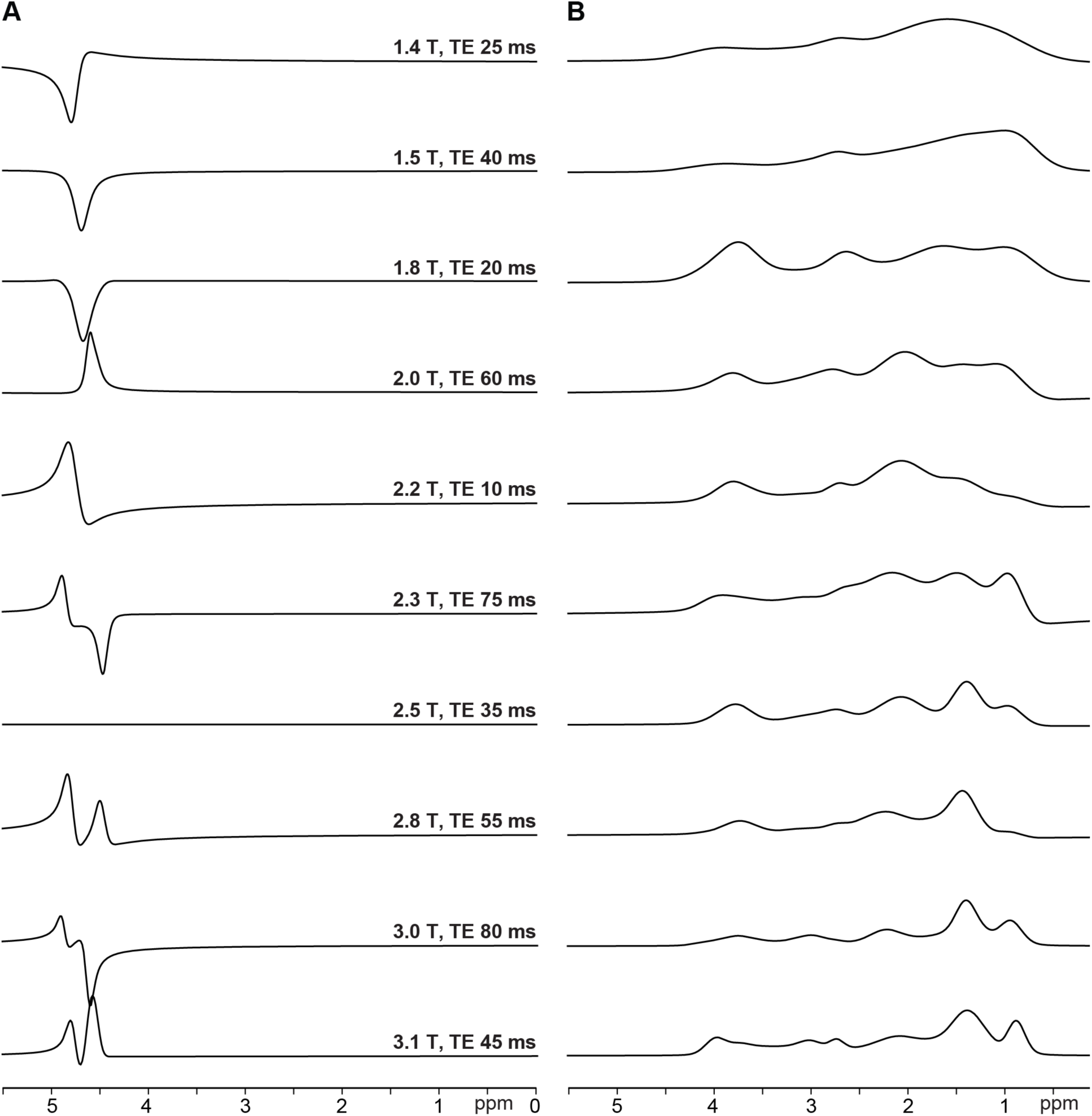
Ten representative examples of **A)** residual water and **B)** macromolecule components. Examples match the full spectra shown in Figure 3; each spectrum is scaled independently for visualization.

**Supplemental Figure 2:**
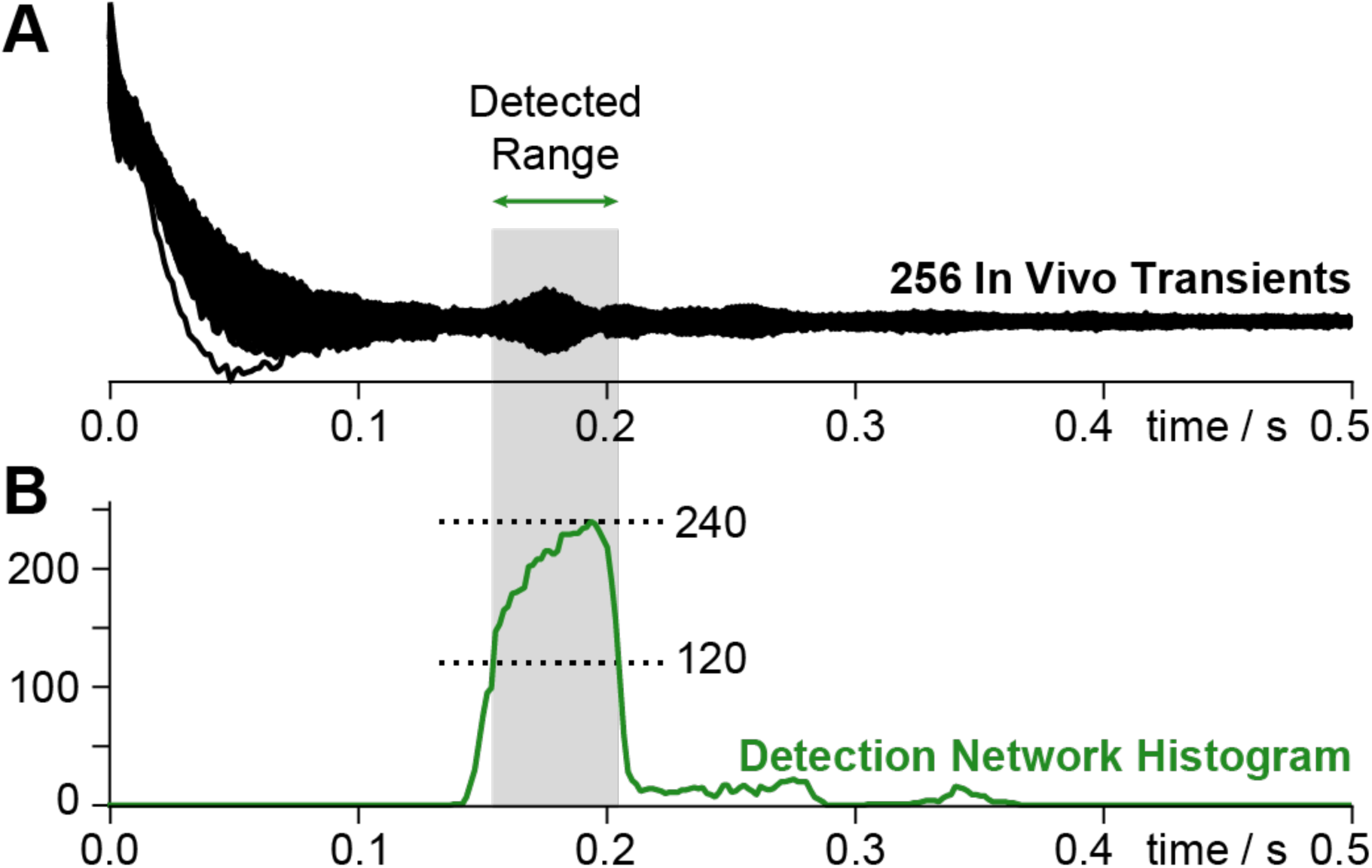
Time-domain window (gray) used to calculate the fractional reduction in standard deviation for the *in vivo* transients. **A)** Each of the 256 MEGA-PRESS transients (128 Edit-on and 128 Edit-Off) overlaid. **B)** Histogram (green) showing the total number of detections by the Detection Network across each timepoint. This window was established algorithmically by using 50% of the maximum count as a threshold for the window.

**Supplemental Table 1:** Concentration ranges for the healthy brain, used to generate synthetic spectra. These mM values were based upon a meta-analysis preliminary to (Gudmundson et al., 2023), with some values (marked *) supplemented from the Fit Challenge ranges (Marjańska et al., 2021) and other ranges (marked †) extended to offer greater flexibility. Concentrations were sampled uniformly between the low and high values to generate the synthetic spectra.

| Metabolite | Synthetic Range (mM) |  | Metabolite | Synthetic Range (mM) |  |
| --- | --- | --- | --- | --- | --- |
|  | Low | High |  | Low | High |
| Acetate | 0.00 | 0.00 | Macromolecule 1.67 | 1.00 | 15.00 |
| Alanine* | 0.47 | 0.77 | Macromolecule 2.04 | 1.00 | 35.00 |
| Ascorbate | 0.36 | 1.53 | Macromolecule 2.26 | 1.00 | 20.00 |
| Aspartate | 0.00 | 4.66 | Macromolecule 2.56 | 1.00 | 5.00 |
| Adenosine Triphosphate | 0.00 | 0.00 | Macromolecule 2.70 | 1.00 | 7.00 |
| β-Hydroxybutyrate | 0.00 | 0.00 | Macromolecule 2.99 | 1.00 | 10.00 |
| β-Hydroxyglutarate | 0.00 | 0.00 | Macromolecule 3.21 | 1.00 | 7.00 |
| Citrate | 0.00 | 0.00 | Macromolecule 3.62 | 1.00 | 5.00 |
| Creatine | 1.41 | 10.50 | Macromolecule 3.75 | 1.00 | 10.00 |
| Cysteine | 0.00 | 0.00 | Macromolecule 3.86 | 1.00 | 4.00 |
| Ethanol Amine | 0.00 | 0.00 | Macromolecule 4.03 | 1.00 | 7.00 |
| Ethyl Alcohol | 0.00 | 0.00 | Myo-inositol<sup>†</sup> | 2.08 | 14.00 |
| γ-Amino Butyric Acid | 0.52 | 1.99 | N-Acetylaspartate<sup>†</sup> | 5.38 | 18.00 |
| Glucose* | 0.94 | 1.53 | N-Acetylaspartylglutamate<sup>†</sup> | 0.26 | 2.26 |
| Glutamine | 0.26 | 3.64 | Phosphocholine* | 0.01 | 2.00 |
| Glutamate | 3.88 | 13.17 | Phosphocreatine* | 3.38 | 6.44 |
| Glycerophosphocholine | 0.05 | 5.00 | Phosphoethanolamine* | 1.41 | 2.30 |
| Glutathione | 0.16 | 2.41 | Phosphoethyl Alcohol | 0.00 | 0.00 |
| Glycine* | 0.94 | 1.53 | Phenylalanine | 0.00 | 0.00 |
| Glycerol | 0.00 | 0.00 | Scyllo-inositol | 0.00 | 0.39 |
| Histamine | 0.00 | 0.00 | Serine | 0.00 | 0.00 |
| Histidine | 0.00 | 0.00 | Taurine | 0.00 | 2.89 |
| Homocarnosine | 0.00 | 0.00 | Threonine | 0.00 | 0.00 |
| Lactate | 0.00 | 1.44 | Tryptophan | 0.00 | 0.00 |
| Macromolecule 0.92 | 1.00 | 30.00 | Tyrosine | 0.00 | 0.00 |
| Macromolecule 1.21 | 1.00 | 8.00 | Valine | 0.00 | 0.00 |
| Macromolecule 1.39 | 1.00 | 35.00 |  |  |  |

**Supplemental Table 2:** Clinical population scaling factors used to generate synthetic spectra. In each case the simulated concentration for a given clinical spectrum was determined by a uniformly sampled concentration drawn from the ranges shown in Supplemental Table 1, multiplied by a scaling factor determined by a uniformly sampled scalar from these ranges provided in Supplemental Table 2.

| Disease / Metabolite | Synthetic Range |  | Metabolite | Synthetic Range |  |
| --- | --- | --- | --- | --- | --- |
|  | Low | High |  | Low | High |
| Seizure |  |  | Cancer |  |  |
| Creatine | 0.918 | 1.012 | Creatine | 0.256 | 1.340 |
| Phosphocreatine | 0.918 | 1.012 | Phosphocreatine | 0.256 | 1.340 |
| Glycerophosphocholine | 0.731 | 1.147 | Glycerophosphocholine | 1.139 | 1.949 |
| Phosphocholine | 0.731 | 1.147 | Phosphocholine | 1.139 | 1.949 |
| γ-Amino Butyric Acid | 0.930 | 1.173 | Glutamate | 0.780 | 1.320 |
| Glutamate | 0.787 | 1.247 | Glutamine | 0.780 | 1.320 |
| Glutamine | 0.787 | 1.247 | Lactate | 1.00 | 9.99 |
| Glutathione | 0.887 | 1.243 | Myo-inositol | 0.829 | 1.519 |
| Myo-inositol | 0.802 | 1.134 | N-Acetylaspartate | 0.509 | 0.956 |
| N-Acetylaspartate | 0.751 | 1.002 | N-Acetylaspartylglutamate | 0.509 | 0.956 |
| N-Acetylaspartylglutamate | 0.751 | 1.002 | Chronic Pain |  |  |
| Stroke |  |  | Glycerophosphocholine | 0.943 | 1.285 |
| Creatine | 0.684 | 1.146 | Phosphocholine | 0.943 | 1.285 |
| Phosphocreatine | 0.684 | 1.146 | γ-Amino Butyric Acid | 0.896 | 1.168 |
| Glycerophosphocholine | 0.855 | 1.527 | Glutamate | 0.790 | 1.121 |
| Phosphocholine | 0.855 | 1.527 | Glutamine | 0.790 | 1.121 |
| Glutamate | 0.874 | 1.140 | Myo-inositol | 0.942 | 1.049 |
| Glutamine | 0.874 | 1.140 | N-Acetylaspartate | 0.775 | 1.280 |
| Lactate | 1.000 | 6.922 | N-Acetylaspartylglutamate | 0.775 | 1.280 |
| Myo-inositol | 0.827 | 1.265 | Migraine |  |  |
| N-Acetylaspartate | 0.727 | 1.074 | Aspartate | 0.434 | 1.409 |
| N-Acetylaspartylglutamate | 0.727 | 1.074 | Creatine | 0.921 | 1.011 |
| Traumatic Brain Injury |  |  | Phosphocreatine | 0.921 | 1.011 |
| Aspartate | 0.785 | 0.910 | Glycerophosphocholine | 0.959 | 1.137 |
| Creatine | 0.814 | 1.162 | Phosphocholine | 0.959 | 1.137 |
| Phosphocreatine | 0.814 | 1.162 | Glutamate | 0.841 | 1.119 |
| Glycerophosphocholine | 0.930 | 1.057 | Glutamine | 0.841 | 1.119 |
| Phosphocholine | 0.930 | 1.057 | Myo-inositol | 0.866 | 1.032 |
| γ-Amino Butyric Acid | 0.860 | 0.984 | N-Acetylaspartate | 0.755 | 1.067 |
| Glutamate | 0.824 | 1.214 | N-Acetylaspartylglutamate | 0.755 | 1.067 |
| Glutamine | 0.824 | 1.214 | Fibromyalgia |  |  |
| Myo-inositol | 0.737 | 1.315 | Creatine | 0.760 | 1.429 |
| N-Acetylaspartate | 0.795 | 1.011 | Phosphocreatine | 0.760 | 1.429 |
| N-Acetylaspartylglutamate | 0.795 | 1.011 | Glycerophosphocholine | 0.840 | 1.236 |
| Type-1 Diabetes |  |  | Phosphocholine | 0.840 | 1.236 |
| Aspartate | 0.895 | 1.496 | γ-Amino Butyric Acid | 0.724 | 0.937 |
| Creatine | 0.977 | 1.039 | Glutamate | 1.005 | 1.104 |
| Phosphocreatine | 0.977 | 1.039 | Glutamine | 0.711 | 1.107 |
| Glycerophosphocholine | 1.034 | 1.140 | Myo-inositol | 0.844 | 1.232 |
| Phosphocholine | 1.034 | 1.140 | N-Acetylaspartate | 0.847 | 1.061 |
| Glutamate | 0.895 | 1.216 | N-Acetylaspartylglutamate | 0.847 | 1.061 |
| Glutamine | 0.956 | 1.353 |  |  |  |
| Glutathione | 0.872 | 1.435 |  |  |  |
| Myo-inositol | 0.893 | 1.092 |  |  |  |
| N-Acetylaspartate | 0.947 | 1.008 |  |  |  |
| N-Acetylaspartylglutamate | 0.947 | 1.008 |  |  |  |
| Scyllo-inositol | 0.501 | 0.992 |  |  |  |
| Taurine | 0.754 | 1.322 |  |  |  |

| Disease / Metabolite | Synthetic Range |  |
| --- | --- | --- |
|  | Low | High |
| Post-Traumatic Stress Disorder |  |  |
| Creatine | 0.940 | 1.235 |
| Phosphocreatine | 0.940 | 1.235 |
| Glycerophosphocholine | 0.843 | 1.284 |
| Phosphocholine | 0.843 | 1.283 |
| γ-Amino Butyric Acid | 0.982 | 1.059 |
| Glutamate | 0.892 | 1.134 |
| Glutamine | 0.892 | 1.134 |
| Myo-inositol | 0.939 | 1.198 |
| N-Acetylaspartate | 0.969 | 1.156 |
| N-Acetylaspartylglutamate | 0.969 | 1.156 |
| Obsessive Compulsive Disorder |  |  |
| Creatine | 0.890 | 1.320 |
| Phosphocreatine | 0.890 | 1.320 |
| Glycerophosphocholine | 0.784 | 1.223 |
| Phosphocholine | 0.784 | 1.223 |
| Glutamate | 0.868 | 1.243 |
| Glutamine | 0.868 | 1.243 |
| Myo-inositol | 0.743 | 1.437 |
| N-Acetylaspartate | 0.846 | 1.100 |
| N-Acetylaspartylglutamate | 0.846 | 1.100 |
| Depression |  |  |
| Creatine | 0.938 | 1.021 |
| Phosphocreatine | 0.938 | 1.021 |
| Glycerophosphocholine | 0.741 | 1.158 |
| Phosphocholine | 0.741 | 1.158 |
| γ-Amino Butyric Acid | 0.769 | 1.400 |
| Glutamate | 0.872 | 1.119 |
| Glutamine | 0.894 | 1.177 |
| Glutathione | 0.822 | 1.082 |
| Myo-inositol | 0.874 | 1.239 |
| N-Acetylaspartate | 0.864 | 1.080 |
| N-Acetylaspartylglutamate | 0.864 | 1.080 |
| Addiction |  |  |
| Creatine | 0.775 | 1.161 |
| Phosphocreatine | 0.755 | 1.161 |
| Glycerophosphocholine | 0.788 | 1.202 |
| Phosphocholine | 0.788 | 1.202 |
| γ-Amino Butyric Acid | 0.669 | 1.289 |
| Glutamate | 0.807 | 1.229 |
| Glutamine | 0.807 | 1.229 |
| Glycine | 0.969 | 1.335 |
| Glutathione | 0.935 | 1.442 |
| Myo-inositol | 0.820 | 1.135 |
| N-Acetylaspartate | 0.761 | 1.195 |
| N-Acetylaspartylglutamate | 0.761 | 1.195 |
|  | Low | High |
| Schizophrenia |  |  |
| Creatine | 0.948 | 1.045 |
| Phosphocreatine | 0.948 | 1.045 |
| Glycerophosphocholine | 0.946 | 1.157 |
| Phosphocholine | 0.946 | 1.157 |
| γ-Amino Butyric Acid | 0.732 | 1.261 |
| Glutamate | 0.857 | 1.164 |
| Glutamine | 0.857 | 1.164 |
| Myo-inositol | 0.806 | 1.239 |
| N-Acetylaspartate | 0.910 | 1.103 |
| N-Acetylaspartylglutamate | 0.910 | 1.103 |
| Psychosis |  |  |
| Creatine | 0.983 | 1.059 |
| Phosphocreatine | 0.983 | 1.059 |
| Glycerophosphocholine | 0.892 | 1.127 |
| Phosphocholine | 0.892 | 1.127 |
| γ-Amino Butyric Acid | 0.725 | 1.176 |
| Glutamate | 0.813 | 1.172 |
| Glutamine | 0.813 | 1.172 |
| Glycine | 1.131 | 1.423 |
| Glutathione | 0.917 | 1.034 |
| Myo-inositol | 0.892 | 1.090 |
| N-Acetylaspartate | 0.910 | 1.048 |
| N-Acetylaspartylglutamate | 0.910 | 1.048 |
| Personality Disorder |  |  |
| Creatine | 0.961 | 1.110 |
| Phosphocreatine | 0.961 | 1.110 |
| Glycerophosphocholine | 0.925 | 1.007 |
| Phosphocholine | 0.925 | 1.007 |
| Glutamate | 0.949 | 1.207 |
| Glutamine | 0.949 | 1.207 |
| Glutathione | 0.917 | 1.034 |
| Myo-inositol | 0.989 | 1.081 |
| N-Acetylaspartate | 0.880 | 0.997 |
| N-Acetylaspartylglutamate | 0.880 | 0.997 |
| Bipolar Disorder |  |  |
| Creatine | 0.900 | 1.061 |
| Phosphocreatine | 0.900 | 1.061 |
| Glycerophosphocholine | 0.854 | 1.269 |
| Phosphocholine | 0.854 | 1.269 |
| Glutamate | 0.907 | 1.115 |
| Glutamine | 0.907 | 1.115 |
| Glutathione | 0.957 | 1.150 |
| Myo-inositol | 0.812 | 1.209 |
| N-Acetylaspartate | 0.863 | 1.109 |
| N-Acetylaspartylglutamate | 0.863 | 1.109 |

| Disease / Metabolite | Synthetic Range |  | Disease / Metabolite | Synthetic Range |  |
| --- | --- | --- | --- | --- | --- |
|  | Low | High |  | Low | High |
| Multiple Sclerosis |  |  | Dementia |  |  |
| Glycerophosphocholine | 0.880 | 1.077 | Ascorbate | 1.132 | 1.231 |
| Phosphocholine | 0.880 | 1.077 | Aspartate | 1.028 | 1.168 |
| γ-Amino Butyric Acid | 0.851 | 1.017 | Creatine | 1.010 | 1.028 |
| Glutamate | 0.887 | 1.030 | Phosphocreatine | 1.010 | 1.028 |
| Glutamine | 0.887 | 1.030 | Glycerophosphocholine | 0.850 | 1.150 |
| Glutathione | 0.844 | 1.069 | Phosphocholine | 0.850 | 1.150 |
| Myo-inositol | 0.892 | 1.078 | γ-Amino Butyric Acid | 0.513 | 1.183 |
| N-Acetylaspartate | 0.924 | 1.044 | Glutamate | 0.771 | 1.139 |
| N-Acetylaspartylglutamate | 0.924 | 1.044 | Glutamine | 0.955 | 1.172 |
| Parkinson's Disease |  |  | Myo-inositol | 0.801 | 1.397 |
| Creatine | 0.850 | 1.100 | N-Acetylaspartate | 0.723 | 1.038 |
| Phosphocreatine | 0.850 | 1.100 | N-Acetylaspartylglutamate | 0.723 | 1.038 |
| Glycerophosphocholine | 0.780 | 1.201 | Scyllo-inositol | 0.476 | 1.312 |
| Phosphocholine | 0.780 | 1.201 | Taurine | 0.882 | 1.013 |
| γ-Amino Butyric Acid | 0.679 | 1.390 | APOE4 | | |
| Glutamate | 0.887 | 1.224 | Aspartate | 1.028 | 1.168 |
| Glutamine | 0.887 | 1.224 | Glycerophosphocholine | 0.965 | 1.019 |
| Myo-inositol | 0.810 | 1.190 | Phosphocholine | 0.965 | 1.019 |
| N-Acetylaspartate | 0.756 | 1.240 | γ-Amino Butyric Acid | 0.513 | 1.183 |
| N-Acetylaspartylglutamate | 0.756 | 1.240 | Glucose | 0.971 | 1.028 |
| Essential Tremor |  |  | Glutamate | 0.836 | 1.126 |
| Creatine | 0.924 | 1.053 | Glutamine | 0.909 | 1.232 |
| Phosphocreatine | 0.924 | 1.053 | Glutathione | 0.834 | 1.103 |
| Glycerophosphocholine | 0.851 | 1.044 | Myo-inositol | 0.959 | 1.092 |
| Phosphocholine | 0.851 | 1.044 | N-Acetylaspartate | 0.895 | 1.063 |
| γ-Amino Butyric Acid | 0.802 | 1.218 | N-Acetylaspartylglutamate | 0.895 | 1.063 |
| Glutamate | 1.050 | 1.434 |  |  |  |
| Glutamine | 1.050 | 1.434 |  |  |  |
| N-Acetylaspartate | 0.919 | 1.136 |  |  |  |
| N-Acetylaspartylglutamate | 0.919 | 1.136 |  |  |  |

**Supplemental Table 3:** T2 Relaxation time ranges in milliseconds for the healthy brain derived from 1.5 T multiple meta-regression preliminary to (Gudmundson et al., 2023). Relaxation times were sampled uniformly between the low and high values.

| Metabolite | Synthetic Range (ms) |  | Metabolite | Synthetic Range (ms) |  |
| --- | --- | --- | --- | --- | --- |
|  | Low | High |  | Low | High |
| Acetate | 0.00 | 0.00 | Macromolecule 1.67 | 20.00 | 60.00 |
| Alanine* | 100.00 | 250.00 | Macromolecule 2.04 | 20.00 | 60.00 |
| Ascorbate | 100.00 | 250.00 | Macromolecule 2.26 | 20.00 | 60.00 |
| Aspartate | 120.15 | 204.55 | Macromolecule 2.56 | 20.00 | 60.00 |
| Adenosine Triphosphate | 0.00 | 0.00 | Macromolecule 2.70 | 20.00 | 60.00 |
| β-Hydroxybutyrate | 0.00 | 0.00 | Macromolecule 2.99 | 20.00 | 60.00 |
| β-Hydroxyglutarate | 0.00 | 0.00 | Macromolecule 3.21 | 20.00 | 60.00 |
| Citrate | 0.00 | 0.00 | Macromolecule 3.62 | 20.00 | 60.00 |
| Creatine 3.03 | 164.08 | 242.70 | Macromolecule 3.75 | 20.00 | 60.00 |
| Creatine 3.91 | 135.18 | 213.80 | Macromolecule 3.86 | 20.00 | 60.00 |
| Creatine 6.65 | 164.08 | 242.70 | Macromolecule 4.03 | 20.00 | 60.00 |
| Cysteine | 0.00 | 0.00 | Myo-inositol<sup>†</sup> | 139.80 | 219.58 |
| Ethanolamine | 0.00 | 0.00 | N-Acetylaspartate | 242.70 | 320.17 |
| Ethyl Alcohol | 0.00 | 0.00 | N-Acetylaspartylglutamate | 132.87 | 216.11 |
| γ-Amino Butyric Acid | 77.37 | 161.77 | Phosphocholine | 100 | 250 |
| Glucose | 100.00 | 250.00 | Phosphocreatine 3.03 | 130 | 210 |
| Glutamine | 103.96 | 184.89 | Phosphocreatine 3.93 | 100 | 180 |
| Glutamate | 140.96 | 219.58 | Phosphocreatine 6.58 | 130 | 210 |
| Glycerophosphocholine | 198.77 | 278.54 | Phosphocreatine 7.30 | 130 | 210 |
| Glutathione | 108.59 | 188.36 | Phosphoethanolamine | 100 | 250 |
| Glycine | 121.31 | 204.55 | Phosphoethyl Alcohol | 0.00 | 0.00 |
| Glycerol | 0.00 | 0.00 | Phenylalanine | 0.00 | 0.00 |
| Histamine | 0.00 | 0.00 | Scyllo-inositol | 100 | 250 |
| Histidine | 0.00 | 0.00 | Serine | 0.00 | 0.00 |
| Homocarnosine | 0.00 | 0.00 | Taurine | 151.37 | 231.14 |
| Lactate | 142.12 | 226.52 | Threonine | 0.00 | 0.00 |
| Macromolecule 0.92 | 20.00 | 60.00 | Tryptophan | 0.00 | 0.00 |
| Macromolecule 1.21 | 20.00 | 60.00 | Tyrosine | 0.00 | 0.00 |
| Macromolecule 1.39 | 20.00 | 60.00 | Valine | 0.00 | 0.00 |

**Supplemental Table 4:** Field strengths (*Tesla*) and possible spectral widths (*Hertz*) available using the AGNOSTIC basis sets. These combinations are achievable by subsampling the time-domain from every timepoint to every 8^th^ timepoint and allows for maintaining a minimum of 2048 timepoints. Each of these combinations is available for the 15 echo times, from 10 ms to 80 ms, in steps of 5 ms.

| **B<sub>0</sub> (T)** | **Every**<br>**Point** | **Every**<br>**2<sup>nd</sup>**<br>**Point** | **Every**<br>**3<sup>rd</sup>**<br>**Point** | **Every**<br>**4<sup>th</sup> Point** | **Every**<br>**5<sup>th</sup> Point** | **Every**<br>**6<sup>th</sup> Point** | **Every**<br>**7<sup>th</sup> Point** | **Every**<br>**8<sup>th</sup> Point** |
| --- | --- | --- | --- | --- | --- | --- | --- | --- |
| 1.4 | 3733.33 | 1866.67 | 1244.44 | 933.33 | 746.67 | 622.22 | 533.33 | 466.67 |
| 1.5 | 4000.00 | 2000.00 | 1333.33 | 1000.00 | 800.00 | 666.67 | 571.43 | 500.00 |
| 1.6 | 4266.67 | 2133.33 | 1422.22 | 1066.67 | 853.33 | 711.11 | 609.52 | 533.33 |
| 1.7 | 4533.33 | 2266.67 | 1511.11 | 1133.33 | 906.67 | 755.56 | 647.62 | 566.67 |
| 1.8 | 4800.00 | 2400.00 | 1600.00 | 1200.00 | 960.00 | 800.00 | 685.71 | 600.00 |
| 1.9 | 5066.67 | 2533.33 | 1688.89 | 1266.67 | 1013.33 | 844.44 | 723.81 | 633.33 |
| 2.0 | 5333.33 | 2666.67 | 1777.78 | 1333.33 | 1066.67 | 888.89 | 761.90 | 666.67 |
| 2.1 | 5600.00 | 2800.00 | 1866.67 | 1400.00 | 1120.00 | 933.33 | 800.00 | 700.00 |
| 2.2 | 5866.67 | 2933.33 | 1955.56 | 1466.67 | 1173.33 | 977.78 | 838.10 | 733.33 |
| 2.3 | 6133.33 | 3066.67 | 2044.44 | 1533.33 | 1226.67 | 1022.22 | 876.19 | 766.67 |
| 2.4 | 6400.00 | 3200.00 | 2133.33 | 1600.00 | 1280.00 | 1066.67 | 914.29 | 800.00 |
| 2.5 | 6666.67 | 3333.33 | 2222.22 | 1666.67 | 1333.33 | 1111.11 | 952.38 | 833.33 |
| 2.6 | 6933.33 | 3466.67 | 2311.11 | 1733.33 | 1386.67 | 1155.56 | 990.48 | 866.67 |
| 2.7 | 7200.00 | 3600.00 | 2400.00 | 1800.00 | 1440.00 | 1200.00 | 1028.57 | 900.00 |
| 2.8 | 7466.67 | 3733.33 | 2488.89 | 1866.67 | 1493.33 | 1244.44 | 1066.67 | 933.33 |
| 2.9 | 7733.33 | 3866.67 | 2577.78 | 1933.33 | 1546.67 | 1288.89 | 1104.76 | 966.67 |
| 3.0 | 8000.00 | 4000.00 | 2666.67 | 2000.00 | 1600.00 | 1333.33 | 1142.86 | 1000.00 |
| 3.1 | 8266.67 | 4133.33 | 2755.56 | 2066.67 | 1653.33 | 1377.78 | 1180.95 | 1033.33 |

